# [^18^F]-Florbetapir PET: Towards Predicting Dementia in Adults with Down Syndrome

**DOI:** 10.1101/235440

**Authors:** David B. Keator, Eric Doran, Theo G.M. Van Erp, Michael J. Phelan, Katherine Tseung, Michael A. Yassa, Steven G. Potkin, Ira T. Lott

## 1. Background

Individuals with Down syndrome (DS) have an increasing age-related prevalence of Alzheimer’s disease (AD). In DS, the triplication of amyloid precursor protein (APP) on chromosome 21 contributes to a life-long accumulation of brain amyloid (Aβ) [1]. Dyshomeostasis of Aβ has emerged as one of the most well-validated factors in the pathogenesis of AD [2]. By age 40 years, virtually all individuals with DS have amyloid plaques and tau tangles, hallmark pathologies that are characteristic of sporadic AD in the general population [3]. Dementia, while not inevitable in DS, increases significantly in prevalence with age to over 75% after age 65 years. [4–6]. Early identification of those who are at the most risk for transitioning to dementia is paramount to intervention trials in DS since improvement is less effective after symptoms of cognitive decline are observable [7].

The urgency to find viable biomarkers for transition to dementia is shared across individuals with DS and those at risk for AD in the general population. A variety of imaging, neuropsychological and genetic biomarkers have been studied to predict conversion to AD in the general population [8]. These approaches have included Positron Emission Tomography (PET) using ligands identifying brain amyloid accumulation ([^18^F]Florbetapir, [^11^C]PiB, and [^18^F]FDDNP) [9–14]. In the general population, the biomarker utility of increased cortical uptake as a result of amyloid binding on PET scans has been of great interest in preclinical AD. However, it is not clear that the uptake data reliably predicts those who will transition to dementia [15,16]. Various working groups in nuclear medicine and AD have developed criteria for using amyloid PET in the diagnosis of patients with persistent or unexplained mild cognitive impairment in the general population but the diagnostic and predictive status of these measures remains uncertain [17]. At the present time, amyloid-PET uptake is best utilized in conjunction with other biomarkers such as CSF Aβ and tau concentrations to assist in the differential diagnosis and prognosis of AD in the general population [18]. At issue is whether amyloid-PET alone may be predictive of dementia in DS.

In DS, [^11^C] PiB binding in individuals with DS appears first in the striatum around 40 years of age and is strongly associated with dementia and cognitive decline [19,20]. [^18^F]-Florbetapir binding in DS reflects fibrillar Aβ, the protein associated with many amyloidoses including AD [21]. Brain amyloid binding as measured by [^18^F]-Florbetapir (Amyvid) PET increases with age in both demented and nondemented participants with DS [22]. The two best predictors of PiB binding in DS are age and a clinical diagnosis of AD [23]. Since Aβ accumulates in the brain of individuals across the lifespan in DS, it may be possible to ascertain a signal in Aβ binding patterns or spatial distribution that predicts a conversion to dementia. The aim of the current study is to evaluate the relationship between baseline [^18^F]-Florbetapir uptake patterns and subsequent transition to dementia in DS. We hypothesized greater amyloid burden in individuals with DS who transition to AD compared to those who do not transition to AD (dementia).

## 2. Methods

### 2.1. Participant Characteristics

Nineteen non-demented adult participants with DS were clinically evaluated at baseline, 9, 18, 27, and 48 months. Informed consent was obtained from each participant prior to enrollment in the study. During the study, five participants transitioned to dementia based on clinical evaluations (supplement S2). The cohort characteristics are shown in Table 1. The groups did not differ in mean age (t(17)=0.70;p<0.49) or gender (p<0.30 Fisher’s) and handedness (p<0.65 Fisher’s) distributions.

**Table 1:** Participant characteristics showing mean (±std) time from PET scan to clinical transition, gender, handedness, and mean (±std) age at time of PET scan.

| Group | N | Time to Transition (yrs.)<br>[min,max] | Gender | Handedness | Age at PET(yrs.)<br>[min,max] |
| --- | --- | --- | --- | --- | --- |
| Non-Transitioned | 14 | NA | 10 Male<br>4 Female | 10 Right<br>3 Left<br>1 Mixed | $50.4 \pm 4.3$<br>[42.8, 58.5] |
| Transitioned | 5 | $1.9 \pm 1.3$<br>[0.6, 3.3] | 2 Male<br>3 Female | 5 Right | $52.1 \pm 5.7$<br>[42.1, 56.1] |

### 2.2. Diagnosis of Transition to Dementia

Dementia was diagnosed in accordance with ICD-10 and DSM-IV-TR as outlined by Sheehan et al. [24]. The clinical diagnosis followed comprehensive baseline and longitudinal assessments including history, neurological examination, and consideration of previous studies in the medical record. Participants were examined by 2 independent neurologists who were blind to the PET scans and neuropsychological assessments. Confounding conditions, such as sensory deficit, untreated thyroid dysfunction, and major depression, that might mimic the symptoms of dementia, were eliminated during the examination process. The diagnosis of transition to dementia was based upon published guidelines for the diagnosis of AD [25]. Details regarding transition symptoms of individual participants are given in the supplemental material (supplement S2). If there was disagreement between the 2 independent neurologists in the diagnosis of transition then the participant remained as non-transitioned.

### 2.3. Neuropsychological Assessments

To evaluate cognitive changes in the DS cohort over time, the Severe Impairment Battery (SIB) [26], Rapid Assessment for Developmental Disabilities (RADD)[27], Fuld Object Memory Evaluation (FULD) [28], Dementia Questionnaire for Persons with Mental Retardation (DMR) Sum of Cognitive Scores (SCS) [29], and DMR Sum of Social Scores (SOS) neuropsychological assessments were collected at baseline, 9, 18, 27, 36, and 48 months on participants completing the study. The SIB assesses social interaction, short-term memory, orientation/attention, language/reading, general knowledge, and praxis/construction. The FULD assesses naming skills, learning, and short-term memory. The RADD assesses language, orientation/attention, short-term memory, general knowledge, arithmetic, and sensorimotor function.

### 2.4. Image Acquisition

[^18^F]-Florbetapir PET scans were acquired on nineteen participants with DS at the University of California, Irvine Neuroscience Imaging Center using the High Resolution Research Tomograph (HRRT, Siemens Medical Systems[30]). Image acquisition followed the Alzheimer’s Disease Neuroimaging Initiative (ADNI) [31,32] protocol consisting of 4×5 minute frames collected 50-70 minutes after injection of the ligand. PET reconstructions were performed using the 3D ordinary Poisson ordered subset expectation maximization (3D OP-OSEM) algorithm[33,34] with 4 iterations and 16 subsets, yielding an image matrix size of 256×256×207 with 1.22 mm isotropic voxels smoothed with a 3mm full-width at half maximum (FWHM) Gaussian kernel. Transmission scans were collected using a 30 mCi Cesium point source and attenuation correction calculated using maximum a-priori transmission reconstruction (MAP-TR[35,36]), total variation regularization[37], and 3D scatter correction algorithms[38]. Structural T1-weighted MPRAGE scans were acquired for each participant on a 3-Tesla Philips Achieva scanner with the following parameters: orientation: sagittal, TR/TE=6.8/3.2ms, flip angle=9°, NEX=1, field of view=27cm^2^, voxel resolution=0.94×0.94×1.20mm, matrix size=288×288×170, SENSE acceleration factor=2.

### 2.5. Image Preprocessing

PET data preprocessing was performed using the SPM12 Statistical Parametric Mapping tool[39] (RRID:SCR_007037). The reconstructed frames were realigned and averaged prior to analysis. The resulting images were co-registered with their respective MRI scans. Volumetric segmentations of the MRI scans were computed with the FreeSurfer (version 6.0) tool (FS6; RRID:SCR_001847)[40]. MRI-derived regions of interest were extracted in the native MRI space from the FS6 Desikan/Killiany atlas[41] segmentations. The PET counts were converted to standardized uptake value ratio (SUVR) units using a composite reference region based on Landau et al.[42]. The composite reference region was made by averaging the means from the FS6-defined cerebellum, pons/brain stem, and eroded subcortical white matter[42]. A report from the ADNI consortium specifies a cutoff for establishing amyloid positivity of 0.79 SUVR units using the composite reference region[43]. Spatial normalization to the Montreal Neurological Institute (MNI) space was performed by registering the MRI scans to the reference tissue probability maps available in SPM12 and applying the resulting deformation to the co-registered PET scans[44]. The resulting image dimensions were 157×189×156 with 1mm isotropic voxels. After registration to MNI space, images were spatially smoothed with an 8mm FWHM Gaussian kernel prior to statistical analyses. Region of interest (ROI) averages of SUVR-scaled PET scan values were computed for each subject based on the FS6-defined Desikan/Killiany atlas regions[41]. These PET ROI averages were used in the Cox regression and classification analyses.

### 2.6. Data Analysis

In the following sections, we describe analyses designed to evaluate the relationship between regional amyloid burden and clinical transition status.

#### 2.6.1. Group Comparison

All voxel analyses were performed using SPM12. A two-sample ANCOVA was performed comparing regional SUVRs between the participants who clinically transitioned to dementia (n=5) with those who did not (n=14). Covariates included age at PET scan, handedness, and gender. Brain masking was performed using an extrinsic mask from a Downs template created using all available MRI scans. Given the small sample size of the clinically-transitioned group (n=5), a simple sensitivity analysis was performed, removing each subject, one at a time, and computing the group differences. The results of the sensitivity analysis were compared with those of the full-group analysis to evaluate whether the group differences were primarily driven by a single subject (supplement S1).

#### 2.6.2. Time-to-Transition Correlation Analyses

To evaluate the relationship between the time-to-transition measurement and the amyloid binding at baseline, a voxel-wise regression was performed in SPM12. The analysis evaluated the relationship between time-to-transition and baseline SUVRs from the five subjects who transitioned to dementia over the study period. Due to the small number of participants, no additional covariates were included in the model.

#### 2.6.3. Cox Regression Analysis

In DS the risk of developing dementia and AD-related neuropathology is significantly elevated as compared to the general population[45]. Because most, if not all, of the non-transitioned subjects are at risk of developing dementia prior to death, it is of interest to know where in the brain amyloid accumulation significantly affected the risk of clinical conversion based on the exact transition times. To evaluate this risk, Cox regression models [46] were built using the R survival package[47] (RRID:SCR_001905). For each ROI in the Desikan/Killiany atlas, a separate regression model was evaluated which included the ROI SUVR average, age at the time of the PET scan, and gender as additional independent variables.

#### 2.6.4. Classification Analysis of Brain Regions Related to Approximate Transition Times

We evaluated the predictive accuracy of using amyloid PET scan data acquired 0.6-3.3 years before the clinical transition to determine which regions are most predictive of transition. To evaluate these questions logistic regression was used to model the probability of clinical transition status based on the amyloid burden. Separate models were evaluated for each FS6 ROI in a 10-fold cross-validation design with a 75/25 percent training/testing split, stratified by groups (i.e. transitioned, non-transitioned). Mean accuracy, area under the receiver operating characteristic curve (ROC), sensitivity, specificity, and positive/negative predictive value measures were computed for each model across the test folds.

## 3. Results

### 3.1. Amyloid PET comparison of transitioned vs. non-transitioned participants

Results from the voxel-wise group analysis of participants who transitioned to dementia (n=5) compared to those who did not (n=14), with a significance threshold of p<0.01 uncorrected (t(14)=2.6) and extent threshold of p<0.05 (k=3600), are shown in Figure 1. Compared with non-transitioned participants, those who transitioned showed increased amyloid binding in the middle, superior, and inferior orbital frontal lobes, middle and superior temporal lobes, middle cingulum, and insula. Both the middle and superior orbito-frontal and the superior temporal lobes survive a more stringent threshold of p<0.001 uncorrected (t(14)=3.8) and extent threshold of p<0.05 (k=1125). The corresponding Cohen’s d effect sizes ranged from 1.4-3.2, indicating the transitioned participants are above the 90th percentile of amyloid binding relative to the non-transitioned participants in this sample. In subcortical regions, we observed higher SUVRs in participants who transitioned to dementia in the basal ganglia (Transitioned=1.02±0.02;Non-Transitioned=0.98±0.07;Cohen’s *d*=0.88), posterior cingulate (Transitioned=1.09±0.05;Non-Transitioned=1.02±0.05;*d*=1.46), and anterior cingulate (Transitioned=1.05±0.02;Non-Transitioned=1.03±0.05;*d*=0.42) yet these differences were non-significant. The regions identified here are areas of increased amyloid burden in the transitioned group yet there is evidence of amyloid positivity across both groups in a variety of brain regions using the cutoff of 0.79 SUVR units for establishing amyloid positivity from the ADNI consortium (supplement S3)[43].

**Figure 1:**
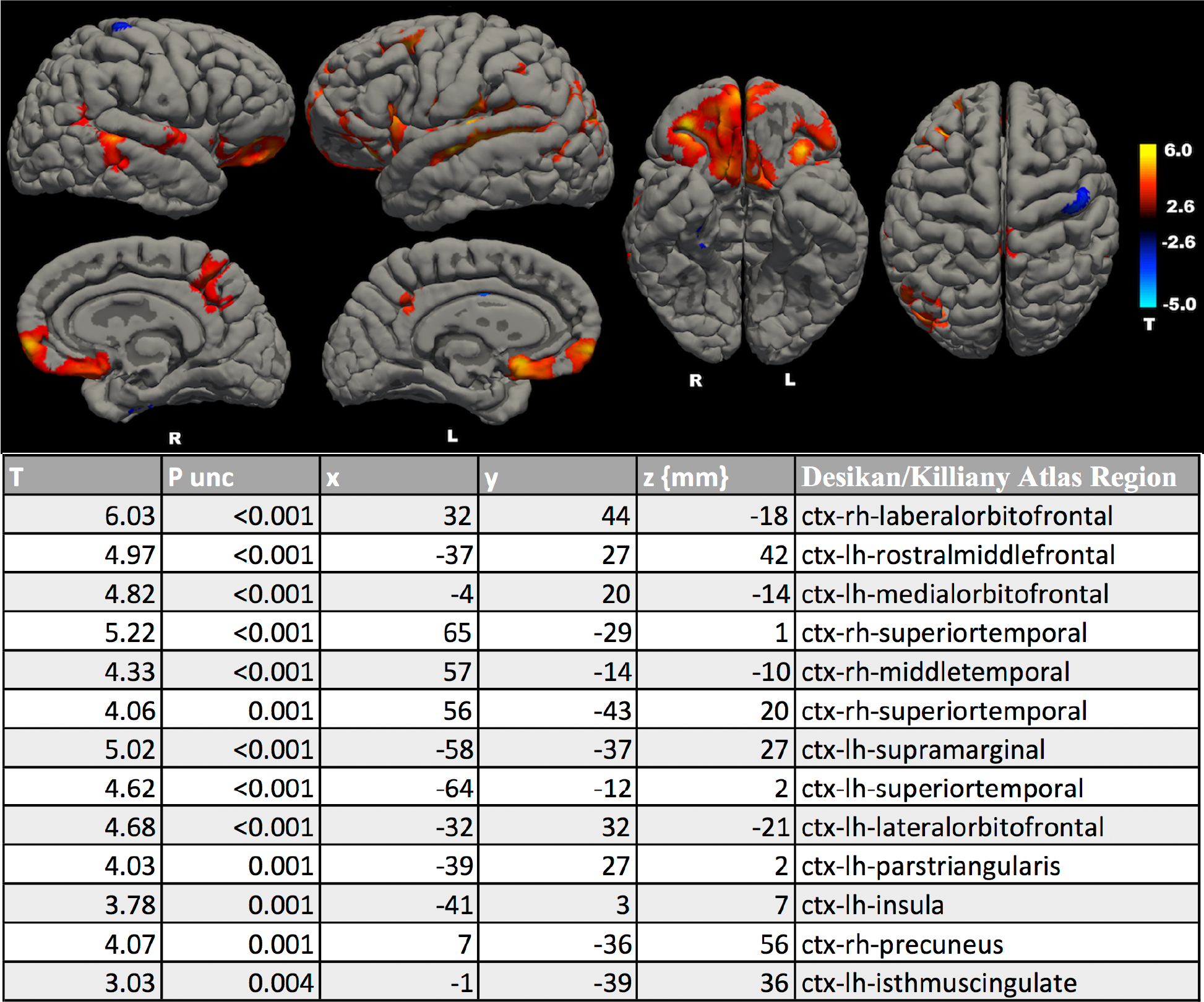
Top: Transitioned vs. non-transitioned group comparison (t(14)= 2.6; p<0.01 uncorrected, p<0.05 extent). Red areas show increased amyloid burden in transitioned subjects. Bottom: Columns contain t-statistic, equivalent z-statistic, uncorrected p-value, x,y, and z MNI coordinates in mm, and the FS6 Desikan/Killiany atlas regions.

### 3.2. Associations between baseline amyloid uptake and transition time

We expect regional amyloid accumulation is related to the state of disease progression. Given the significant variability in transition times (0.6-3.3 years), we computed the voxel-wise association between the amyloid burden at baseline and the transition time after PET acquisition in the five subjects who transitioned. The results are shown in Figure 2 with a significance threshold of p<0.01 uncorrected (t(3)=4.5) and extent threshold of p<0.05 (k=1125). Areas of significant negative correlations (shown in red), corresponding to increased amyloid burden in subjects who transitioned temporally closer to the baseline PET scan, were found in middle and superior orbitofrontal, middle frontal lobe, inferior, middle, and superior temporal lobes, supramarginal, and superior motor area. The middle frontal, superior orbito-frontal, and middle and inferior temporal lobes survive a more stringent threshold of p<0.001 uncorrected (t(3)=10.2) and extent threshold of p<0.05 (k=95). The coefficient of determination (r squared) measures ranged from 0.87-0.98 corresponding to a strong relationship between amyloid burden and the time to transition to clinical dementia.

**Figure 2:**
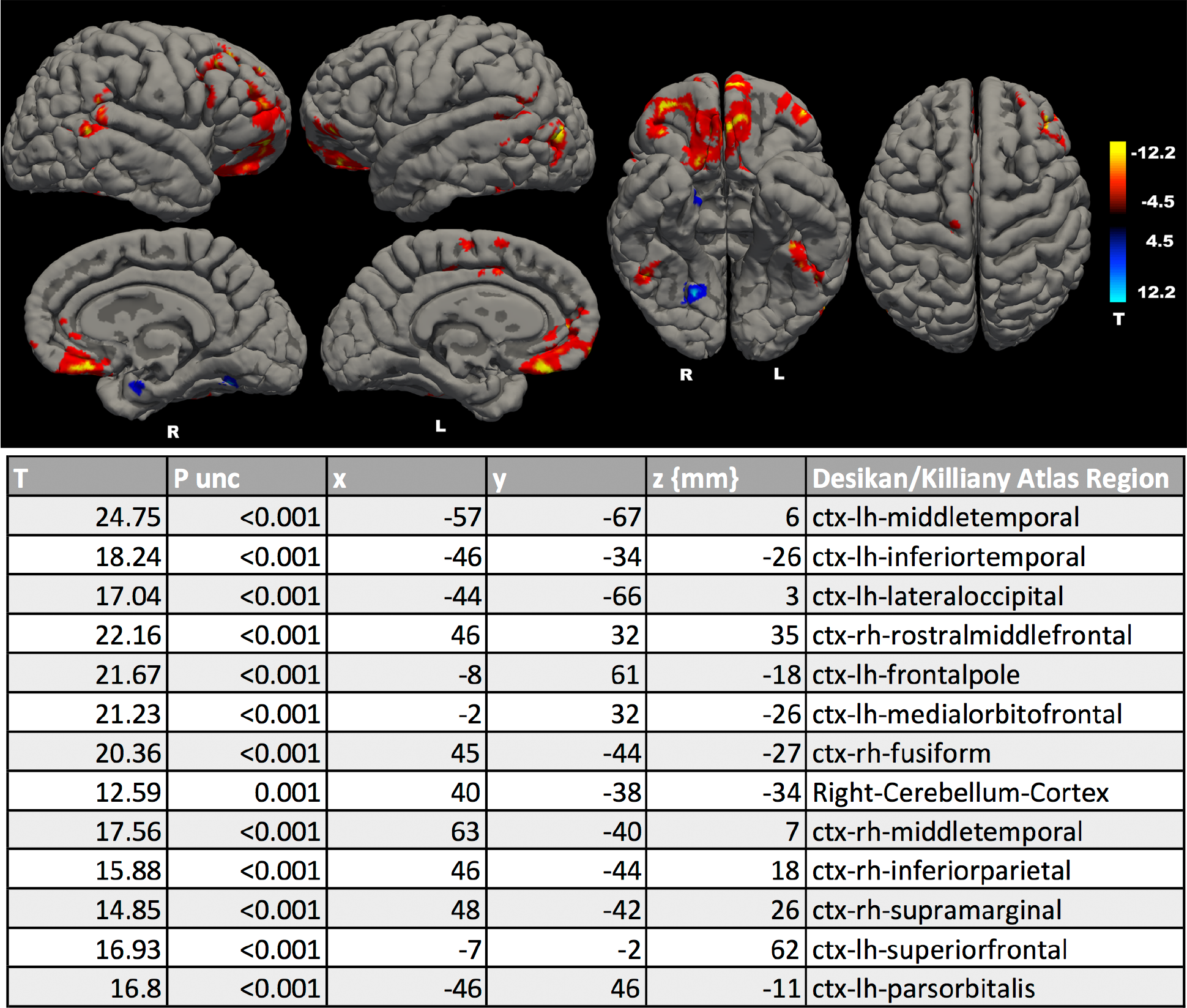
Top: Time-to-transition correlations with amyloid burden (t(3)=4.5; p<0.01 uncorrected, p<0.05 extent). Areas of red correspond to increased amyloid accumulation as the time-to-transition was closer to the baseline PET scan date. Bottom: Columns contain t-statistic, equivalent z-statistic, uncorrected p-value, x, y, and z MNI coordinates in mm, and the FS6 Desikan/Killiany atlas regions.

### 3.3. 3.3. Rate of transition as a function of baseline amyloid burden: Cox Regression Analysis

In evaluating the results of the group analysis and the time-to-transition correlations, we observed areas of significant variability in amyloid binding as a function of transition status and transition time. To understand the influence of the regional amyloid binding on the hazard rate at which the subjects are transitioning, we performed Cox regression analyses for each SUVR ROI average in the FS6 Desikan/Killiany atlas. The top most significant results from the exploratory analysis are shown in Table 2 and consist of regions typically associated with middle to late stages of Alzheimer’s disease neuropathology as described by Braak and Braak[50,51].

**Table 2:**
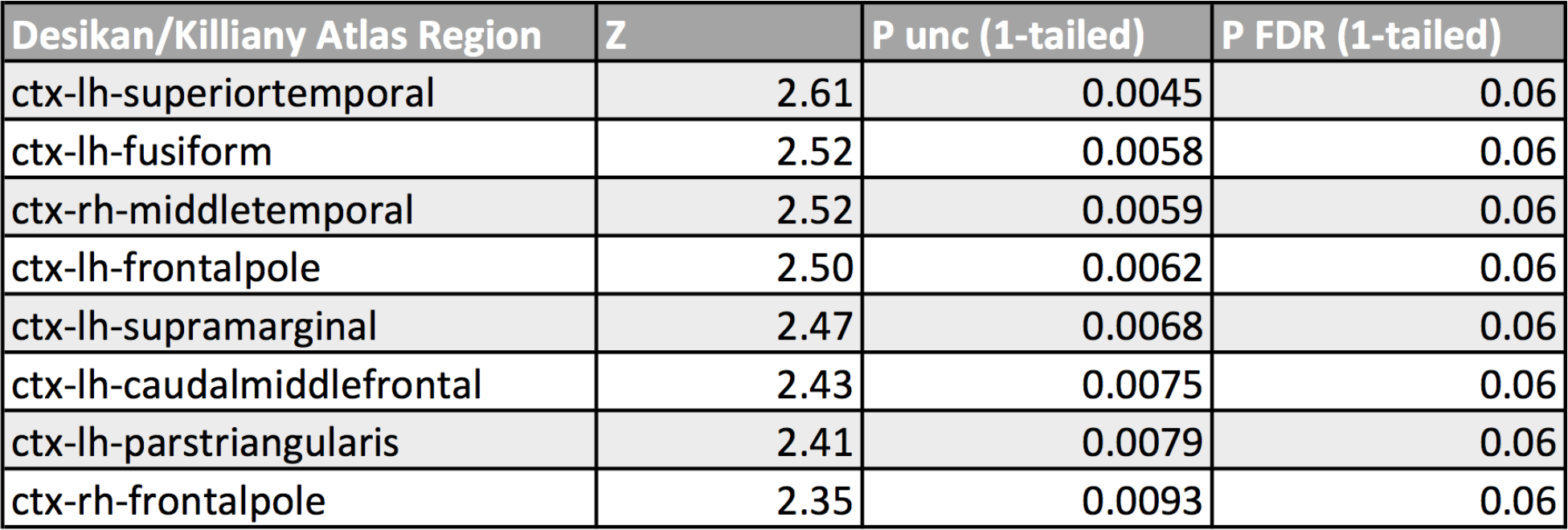
Cox regression analysis results. Columns contain Z, uncorrected one-tailed p-values, and FDR-corrected p-values for FreeSurfer Desikan/Killiany atlas regions with most significant associations.

| Desikan/Killiany Atlas Region | Z | P unc (1-tailed) | P FDR (1-tailed) |
| --- | --- | --- | --- |
| ctx-lh-superiortemporal | 2.61 | 0.0045 | 0.06 |
| ctx-lh-fusiform | 2.52 | 0.0058 | 0.06 |
| ctx-rh-middletemporal | 2.52 | 0.0059 | 0.06 |
| ctx-lh-frontalpole | 2.50 | 0.0062 | 0.06 |
| ctx-lh-supramarginal | 2.47 | 0.0068 | 0.06 |
| ctx-lh-caudalmiddlefrontal | 2.43 | 0.0075 | 0.06 |
| ctx-lh-parstriangularis | 2.41 | 0.0079 | 0.06 |
| ctx-rh-frontalpole | 2.35 | 0.0093 | 0.06 |

### 3.4. Classification using baseline PET ROI averages for predicting future clinical transition

Drawing upon the results from the previous analyses, it was of interest to understand whether regional amyloid burden is predictive of transition. To evaluate this question, we modeled the logit of the probability of transition as a linear function of average amyloid burden for each ROI. The top most accurate results in terms of area under the ROC curve (AUC), sensitivity (Sense) in predicting true positives, specificity (Spec) in predicting true negatives, and positive and negative predictive values (PPV, NPV) in predicting transition status are shown in Table 3. Positive and negative predictive values (PPV, NPV) are the proportions of true positive and true negative predictions from each classifier, relative to the total number of positive and negative predictions respectively. The results are averaged across test cases from all ten folds of cross-validation. To compute results from the trained LR classifiers, the classifiers are evaluated on the test cases from their respective fold using a cutoff probability greater than 0.5 to label a case as transitioned. To compute AUC, the cutoff probability is varied from 0 to 1 and the area under the resulting ROC curve is computed. None of the regions identified differed significantly in anatomical volume between the transitioned and non-transitioned groups, suggesting that these results are unlikely to be confounded by group differences in anatomical structure sizes. The AUC and PPV results suggest that ROI average PET SUVR measurements in regions related to dementia progression may be good predictors of future transition to dementia. For comparison, using the whole-brain average SUVR counts, instead of regionally-specific SUVR averages, resulted in poor classification performance (AUC 0.64±0.20).

**Table 3:** Logistic regression classification analysis results. Columns include mean±standard deviations of: the area under the ROC curves (AUC(Mean±Std), sensitivity and specificity (Sense(Mean±Std), Spec(Mean±Std), positive predictive value (PPV(Mean±Std), and negative predictive values (NPV(Mean±Std) from 10-fold cross validation tests of FS6 Desikan/Killiany atlas regions.

| Desikan/Killiany Atlas Region | AUC(Mean±Std) | Acc(Mean±Std) | Sense (Mean±Std) | Spec (Mean±Std) | PPV(Mean±Std) | NPV(Mean±Std) |
| --- | --- | --- | --- | --- | --- | --- |
| ctx.lh.inferiorparietal | 0.96±0.04 | 0.84±0.12 | 0.92±0.09 | 0.65±0.47 | 0.98±0.04 | 0.75±0.23 |
| ctx.lh.superiortemporal | 0.93±0.10 | 0.87±0.08 | 0.94±0.08 | 0.75±0.27 | 0.90±0.11 | 0.80±0.27 |
| ctx.lh.lateralorbitofrontal | 0.92±0.11 | 0.79±0.15 | 0.85±0.20 | 0.73±0.31 | 0.89±0.13 | 0.74±0.35 |
| ctx.lh.inferiortemporal | 0.90±0.11 | 0.84±0.12 | 0.91±0.13 | 0.77±0.32 | 0.91±0.14 | 0.78±0.31 |
| ctx.rh.precuneus | 0.89±0.09 | 0.80±0.11 | 0.86±0.13 | 0.72±0.30 | 0.88±0.13 | 0.66±0.27 |
| ctx.rh.bankssts | 0.88±0.14 | 0.78±0.13 | 0.90±0.13 | 0.55±0.38 | 0.91±0.09 | 0.71±0.32 |
| ctx.rh.entorhinal | 0.88±0.10 | 0.73±0.08 | 0.82±0.20 | 0.59±0.38 | 0.87±0.14 | 0.60±0.32 |
| ctx.lh.medialorbitofrontal | 0.88±0.15 | 0.80±0.12 | 0.88±0.16 | 0.68±0.31 | 0.88±0.13 | 0.75±0.33 |

### 3.5. Changes in neuropsychological measures over time associated with clinical transition

The participants who transitioned to dementia during the longitudinal observation period showed progressive memory problems, disturbance of gait, decline in their baseline level for carrying out activities of daily living, and decreased productivity in their workshop and community settings (supplement S2). The mean rates of decline and the differences between the transitioned and non-transitioned groups in the neuropsychological assessments are shown in Table 4 and are supportive of the clinical diagnoses of dementia in the group that transitioned. For the RADD, SIB, and FULD assessments, a negative rate of change indicates a worsening of performance. For the DMR SCS and DMR SOS a positive rate of change indicates a worsening of performance (supplement S4). The difference between the average rates of decline on the SIB, for example, in the transitioned group was estimated to be 5.05±1.25 points per year faster than in the non-transition group. Differences in the other assessments can be interpreted in a similar fashion.

**Table 4:** Mean (± standard error) longitudinal rates of change per year in neuropsychological assessments. The transitioned group shows worsening of performance, on average, over time across all assessments.

| Diagnosis | N | RADD | SIB | FULD | DMR-SCS | DMR-SOS |
| --- | --- | --- | --- | --- | --- | --- |
| Non-Transition (NT) | 14 | 0.62±0.59 | 1.23±0.46 | 1.66±1.54 | -0.51±0.28 | -0.27±0.52 |
| Transitioned (T) | 5 | -2.64±0.69 | -3.82±1.16 | -2.95±1.12 | 2.18±1.33 | 2.23±0.87 |
| Difference (NT-T) | | 3.26±0.91 | 5.05±1.25 | 4.61±1.91 | -2.69±1.36 | -2.51±1.02 |

## 4. Discussion

The present work studied a cohort of nineteen participants with DS, five of whom had transitioned to dementia. Transition status was determined by two independent neurologists, blinded to both the baseline PET and neuropsychological assessments. The transitioned cases thus presented a unique opportunity for comparison with those that did not.

A primary goal of the study was to compare the regional distribution of amyloid aggregation on baseline PET images between individuals with DS who did and did not transition to dementia. These analyses described a retrospective case-control study in which mean differences in amyloid burden were quantified across brain regions of interest and related to the clinical diagnosis of dementia. A measure of global amyloid binding was not sensitive to differences between transitioned and non-transitioned participants, as opposed to regional differences which were sensitive and reported in terms of the Cohen’s effect-size d-statistic. Further, exploiting an advantage of logistic regression with retrospective study designs[52], these analyses described the predictive potential for classifying cases based on regional amyloid distribution in baseline scans. Our results suggest that regional amyloid load is informative in evaluating the risk for dementia in DS and, pending replication, is likely a useful quantitative measure to include in a composite risk score.

A second goal of the study was to assess overall decline in support of the clinical evaluations by independent neurologists. Cognitive evaluations were made for all participants at baseline and followed-up prospectively at 9, 18, 27, 36, and 48 months. The rate of change of standard neuropsychological scores was determined from baseline to the month of last follow-up. The results showed that transition status independently tracked trends in RADD, SIB, FULD, and DMR SCS and SOS. Overall, transitioned participants declined faster than non-transitioned participants. These results quantified the rapidity of decline to be expected prospectively of transitioned cases on standard neuropsychological assessments.

The diagnosis of dementia in DS must take into account the variability in baseline intellectual functioning intrinsically associated with trisomy 21. Over time, the clinician’s diagnosis of dementia in DS has been found to be more accurate than the formal criteria put forth in ICD 10 and DSM IV-TR classifications[24,53]. In this report, clinical evaluation by examiners, blinded to the neuropsychological and imaging data, contributed most to the consensus diagnosis of transition status. The data from the neuropsychological battery supported the clinicians’ observations of transition status. In prior studies, the neuropsychological battery used was found to be correlated with clinical dementia status[54,55]. In this study, the slope of the neuropsychological measures correlated with clinical transition status.

Taken together, these results further suggest a role for regional amyloid and neuropsychological assessment—together with tau, and other biomarkers—in developing a risk score for predicting the progression to dementia in Down syndrome. The neuropsychological data are consistent with observations from independent clinical judgement. Clinical judgement plays an important role in the diagnosis of dementia in DS as reinforced by the clinical vignettes on each of the transitioned cases (supplement S2). The rate of β-amyloid accumulation in brains from individuals with DS has been found to differ according to the pre-existing amyloid burden[21,22,58]. Age-related brain atrophy is associated with the rate of PiB-binding and the level of cognitive impairment in DS[59]. Yet amyloid binding after a certain level of retention may not increase with aging in DS[20]. The presence of cortical PiB-binding as a function of age has also been confirmed in post-mortem studies[20,60]. In a study examining global PiB binding as a continuous variable, many adults with DS appeared to tolerate amyloid-β-deposition without deleterious effect on cognitive functioning[53]. While global PiB binding may not predict transition to dementia, our study suggests that there are regional differences in amyloid that may predict cognitive decline.

## 5. Conclusions

This study suggests that [^18^F]-Florbetapir PET scans may be predictive of future clinical transition to dementia in individuals with DS. Future longitudinal studies with larger cohorts of participants will be needed to replicate and validate these findings and to show trajectories of change over time.

## Acknowledgments and Funding

This work was supported in part by NICHD 065160 (Lott), R01 AG053555 (May), and P50 AG16573 (May).

